# Trophectoderm differentiation to invasive syncytiotrophoblast is induced by endometrial epithelial cells during human embryo implantation

**DOI:** 10.1101/2020.10.02.323659

**Authors:** Peter T Ruane, Terence Garner, Lydia Parsons, Phoebe A Babbington, Susan J Kimber, Adam Stevens, Melissa Westwood, Daniel R Brison, John D Aplin

## Abstract

At implantation, trophoblast derived from the trophectoderm of the blastocyst-stage embryo invades the endometrium to establish pregnancy. To understand how embryos breach the endometrial epithelium, we modelled human implantation using blastocysts or trophoblast stem cell spheroids cultured with endometrial epithelial cells (EEC). Blastocyst invasion of the EEC layer was initiated by multinuclear syncytiotrophoblast. Spheroids also invaded the epithelium with syncytiotrophoblast, and EEC induced upregulation of syncytiotrophoblast markers. Modelling implantation *in silico* using blastocyst and EEC transcriptomes revealed gene networks that exhibited greater connectivity and organisation in trophectoderm of the polar region of the embryonic axis. However, gene ontologies and machine learning suggested that EEC drives syncytiotrophoblast differentiation in polar and mural trophectoderm. This is the first evidence for endometrial epithelium-induced trophectoderm differentiation to invasive syncytiotrophoblast as the cellular mechanism of embryonic breaching of the endometrium in humans, with implications for reproductive medicine and our understanding of human embryonic development.

## Introduction

Pregnancy is established as the embryo implants into the endometrium, whereupon placentation enables fetal development. Trophectoderm (TE) forms the outer cell layer of the blastocyst-stage embryo, attaching to the receptive luminal epithelium of the endometrium to initiate implantation before TE-derived trophoblast penetrates the epithelium and invades the endometrial stroma (Aplin & Ruane 2017). Recent progress with *in vitro* culture has revealed details of the cellular choreography of peri-implantation human embryos, up to day 14, in isolation (Deglincerti *et al*. 2016, Shahbazi *et al*. 2016, Shahbazi *et al*. 2017, Popovic *et al*. 2019, West *et al*. 2019, Xiang *et al*. 2019, Zhou *et al*. 2019), and the effects of maternal cell interactions on peri-implantation development up to day 10 have been explored using endometrial stromal cells (Lv *et al*. 2019). However, breaching of the endometrial epithelium in the first phase of human embryo implantation remains largely uncharacterised.

Human embryo implantation *in situ* has been observed only from the early stromal invasion phase in historic samples (Hertig *et al*. 1956). Many studies have reported interaction of the human blastocyst with endometrial epithelial cells (EEC) *in vitro* (Lindenberg *et al*. 1985, Bentin-Ley *et al*. 2000, Galan *et al*. 2000, Meseguer *et al*. 2001, Petersen *et al*. 2005, Lalitkumar *et al*. 2007, Lalitkumar *et al*. 2013, Kang *et al*. 2014, Berger *et al*. 2015, Boggavarapu *et al*. 2016, Aberkane *et al*. 2018, Ruane *et al*. 2020), however, these have predominantly focussed on blastocyst attachment with limited analysis of cellular morphology. Ultrastructural analysis of six blastocysts attached to EEC *in vitro* demonstrated shared desmosomal junctions mediating TE-EEC attachment, TE invasion between EEC, and bi-nucleated TE cells potentially representing syncytiotrophoblast (STB) (Bentin-Ley *et al*. 2000). Trophoblast develops from TE during implantation, and STB forms by cell fusion to produce multinucleated cells. STB was seen to mediate early stromal invasion at human implantation *in vivo* (Hertig *et al*. 1956), and in earlier implantation samples in non-human primates it was observed to penetrate the endometrial epithelium (Enders *et al*. 1983, Smith *et al*. 1987). STB has been seen in human embryos developing beyond the blastocyst stage in isolation (Deglincerti *et al*. 2016, Shahbazi *et al*. 2016, West *et al*. 2019) and in culture with endometrial stromal cells (Lv *et al*. 2019), but whether it mediates embryo breaching of the endometrial epithelium in human remains unknown. Our previous study in mouse embryos suggested that interaction with EEC stimulates trophoblast differentiation (Ruane *et al*. 2017), implicating the epithelial phase of implantation as important for trophoblast development.

The maternal-facing cells of the placenta derive from the TE, and as such a subpopulation of trophoblast must retain multipotency during implantation (Knofler *et al*. 2019), while post-mitotic trophoblast invades the endometrium (Chuprin *et al*. 2013, Lu *et al*. 2017, Velicky *et al*. 2018). Multipotent human trophoblast stem cells (TSC) have recently been derived from blastocysts and first trimester placenta (Okae *et al*. 2018), and from naïve embryonic stem cells (Dong *et al*. 2020), providing promising systems to study implantation. In mice, the pioneering invasive trophoblast derives from the subset of the TE not in contact with the inner cell mass (ICM) of the blastocyst, termed mural TE, while the ICM-adjacent TE, termed polar TE, gives rise to multipotent trophoblast of the ectoplacental cone (Sutherland 2003). *In vitro* studies have demonstrated that human blastocysts orient such that polar TE may mediate attachment to EEC or adhesive surfaces (Bentin-Ley *et al*. 2000, Deglincerti *et al*. 2016, Shahbazi *et al*. 2016, Aberkane *et al*. 2018), while it is not clear from specimens of later stages of human implantation *in vivo* whether polar TE initiates invasion into the endometrium (Hertig *et al*. 1956). Differences between human polar and mural TE have been described at the transcriptomic level (Petropoulos *et al*. 2016), but little is understood about differential function of these TE subpopulations.

We and others have recently characterised human blastocyst attachment to the endometrial epithelial Ishikawa cell line (Nishida *et al*. 1985). This revealed that clinical embryology grading of TE morphology, and blastocyst hatching from the zona pellucida, affects attachment, while adhesion-related genes are upregulated in attached embryos (Aberkane *et al*. 2018, Ruane *et al*. 2020). Here, we further interrogate embryo-EEC interactions *in vitro* and show that STB mediates embryonic breaching of the epithelial layer. Moreover, our TSC spheroid and *in silico* models indicate that attachment to EEC induces STB differentiation.

## Results

### Gross and cellular morphology of embryos attached to Ishikawa EEC layers

Forty six blastocysts attached to Ishikawa EEC layers were analysed in this study, all after co-culture for 48h, from day 6 post-fertilisation to day 8. Attached embryos exhibited either expanded (18/46, 39.1%) (Figure 1A), or collapsed (28/46, 60.9%) blastocyst morphology (Figure 1B). We were able to ascertain whether embryos attached in a the polar TE- or mural TE-oriented fashion in seven instances, with 5/7 exhibiting polar TE-oriented attachment. Phalloidin labelling of cortical actin filaments combined with DAPI labelling of nuclei enabled single cell resolution of embryos attached to EEC layers. Attached embryos with no clear breaching of the epithelium were observed (Figure 1C). Embryos in which a defined region of multinuclear STB was mediating initial breaching of the epithelium and contacting the underlying substrate were also seen (Figure 1D, E). In addition, STB was observed in contact with the apical EEC surface (Figure 1E). The majority of embryos had progressed beyond these initial stages to invade laterally into the EEC layer, with morphologically distinct mononuclear and multinuclear outgrowths present (Figure 1F, G, respectively).

**Figure 1.**
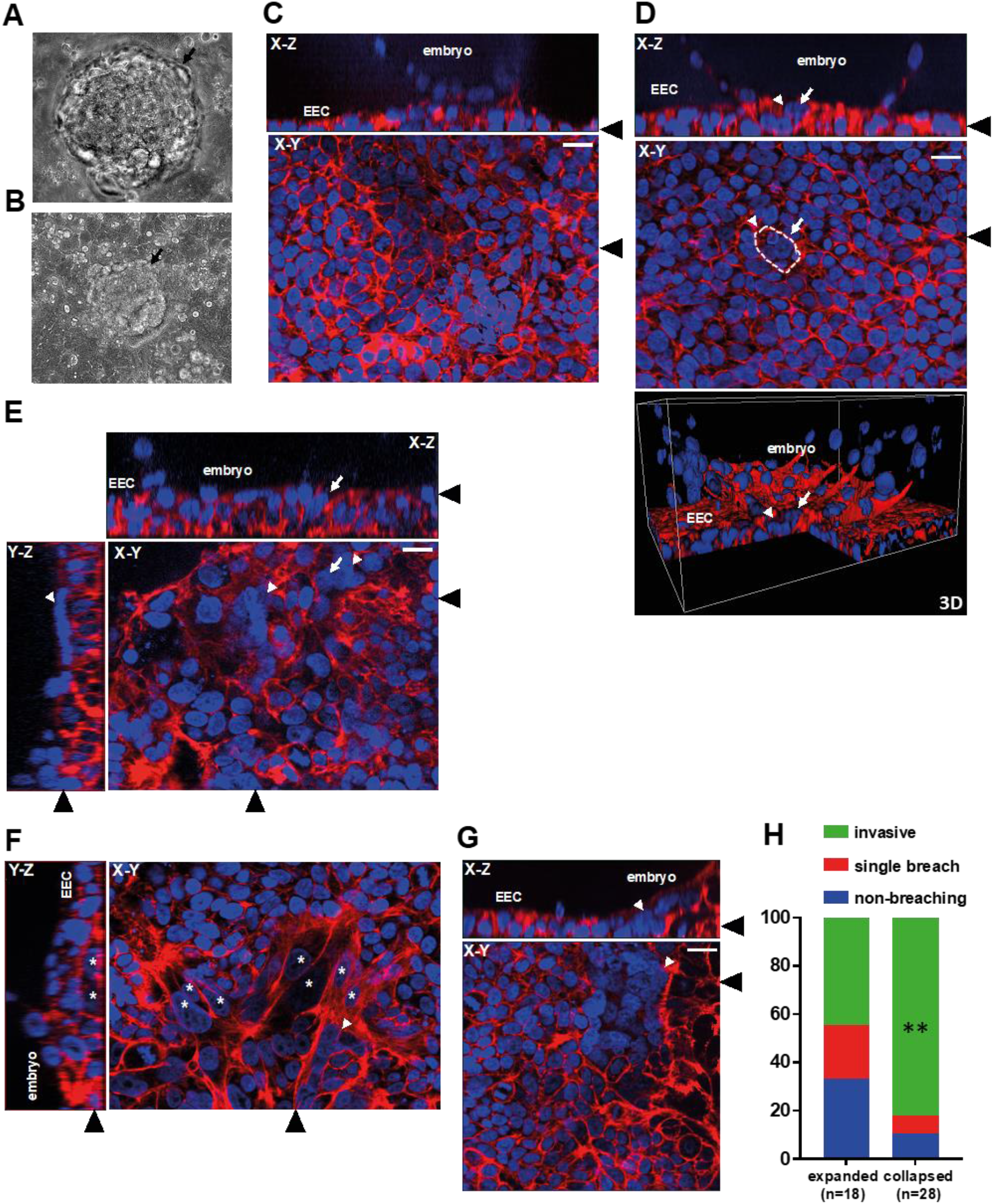
***A, B*** Phase contrast images of embryos attached to Ishikawa EEC layers, showing expanded blastocyst and collapsed blastocyst morphology. Black arrows indicate embryos. Scale bars 20μm. ***C-G*** Attached embryos were stained with phalloidin (red) and DAPI (blue) to label actin and nuclei, respectively, and imaged by optical sectioning fluorescence microscopy. Panels show X-Y, X-Z and Y-Z planes and a 3D image, as labelled. Ishikawa EEC and embryos are indicated in X-Z and Y-Z planes and the 3D image. Black arrowheads indicate the region of the section for the adjoining plane. White arrows indicate trophoblast breaching of the Ishikawa EEC layer, white arrowheads indicate STB, and white asterisks indicate invasive mononuclear trophoblast. Dotted line outlines breaching STB. Scale bars 20μm. ***H*** Bar graph showing proportions of embryo morphologies (n=46). ** p<0.01 Chi-squared

It was observed that 10/18 embryos with expanded blastocyst morphology (55.5%) were superficially attached to the EEC layer or displayed only pioneering breaching of the epithelium, while 8/18 (44.5%) had invaded laterally into the epithelium. In comparison, 5/28 collapsed embryos (17.8%) had not breached or exhibited only pioneering breaching, with 23/28 (82.2%) found to be invasive (Figure 1H). Collapsed embryos were therefore more likely to have extensively invaded the EEC layer. Importantly, each of the seven pioneering breaching events, observed in six embryos, consisted of STB protruding through the EEC layer (Figure 1D, 1E, 2B).

**Figure 2.**
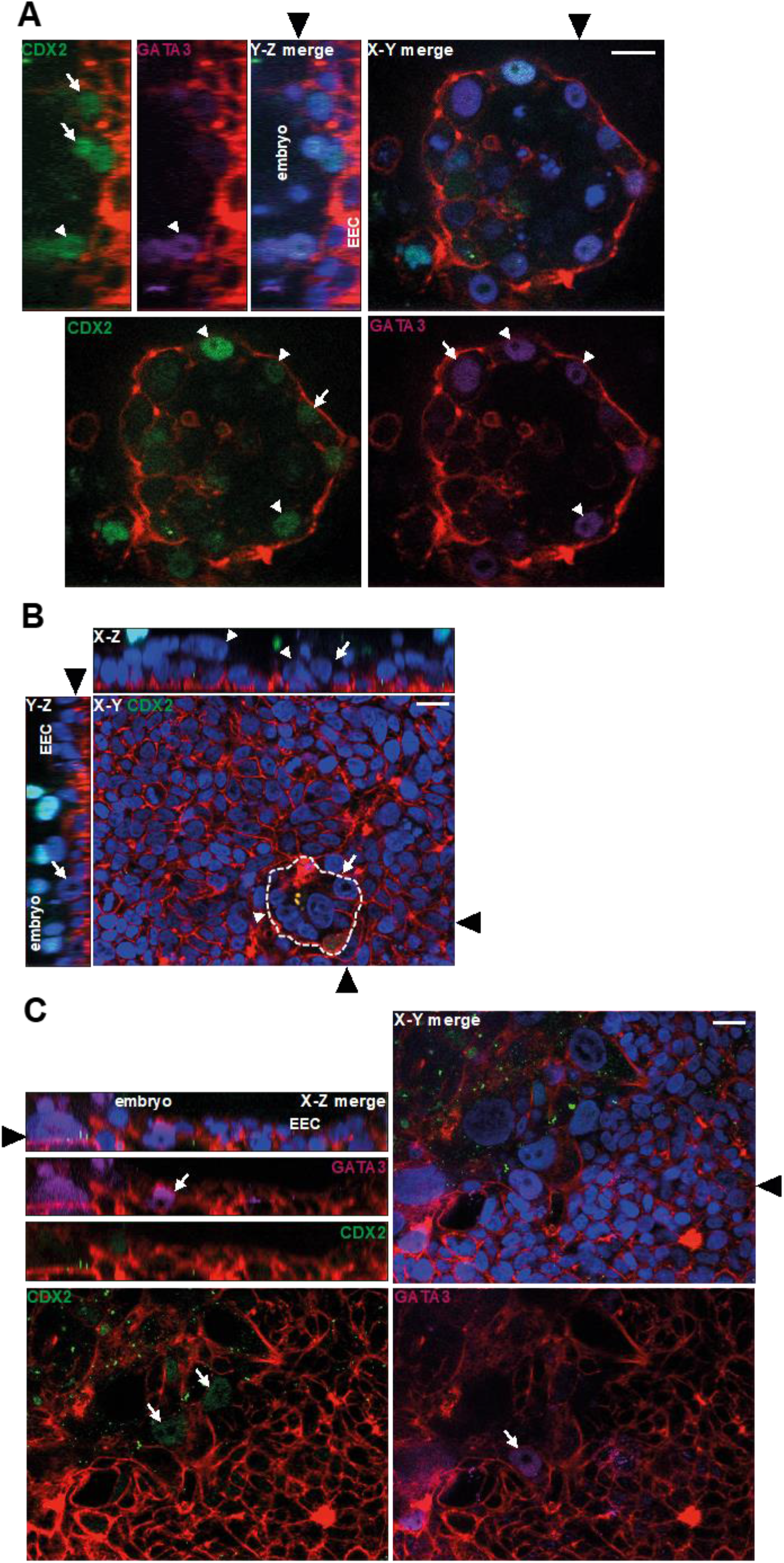
***A*** Fluorescence optical sections of an attached embryo labelled with phalloidin (red), DAPI (blue), anti-CDX2 (green) and anti-GATA3 antibody (magenta). Y-Z and X-Y planes are shown, and Ishikawa EEC and embryos are indicated Y-Z panel. Black arrowheads indicate the location of the adjoining section. White arrows show CDX2 or GATA3-positive TE and white arrowheads indicate double-positive CDX2-GATA3-positive TE. Scale bar 20μm. ***B*** An attached embryo labelled with phalloidin (red), DAPI (blue) and anti-CDX2 antibody (green) and imaged by optical sectioning fluorescence microscopy. The planes shown in each panel are indicated (X-Z, X-Y) and black arrowheads indicate the location of the adjoining section. White arrows point to trophoblast breaching of the Ishikawa EEC layer and white arrowheads indicate STB. Dotted line outlines breaching STB. Scale bar 20μm. ***C*** An invasive embryo labelled with phalloidin (red), DAPI (blue), anti-CDX2 (green) and anti-GATA3 antibody (magenta), optical planes indicated on the panels. White arrows indicate CDX2- or GATA3-positive mononuclear trophoblast. Black arrowheads indicate the location of the adjoining section. Scale bar 20μm.

### TE markers in EEC-attached embryos

To characterise the TE/trophoblast cell types in attached embryos, we labelled samples with antibodies to the transcription factors CDX2 and GATA3, which are expressed in this lineage (Hemberger *et al*. 2020). Co-staining for CDX2 and GATA3 in expanded blastocysts which had not breached the epithelium revealed heterogeneity amongst TE both in contact with, and distal to, EEC; some nuclei were positive only for CDX2, some only for GATA3 and yet others co-labelled with both (Figure 2A). In embryos which had just breached the epithelium, CDX2 was present only in embryonic nuclei above the plane of the EEC layer (Figure 2B). In collapsed embryos, CDX2 and GATA3 labelling was present in invasive mononuclear trophoblast (Figure 2C). Neither transcription factor was detected in STB.

### Multiple STB regions invade into the EEC layer

E-cadherin is present at intercellular junctions in human TE and mononuclear trophoblasts (Alikani 2005, Aplin *et al*. 2009), and was used to demarcate mononuclear cells. Multinuclear elements lacking inter-nuclear E-cadherin labelling were observed in contact with EEC at the invading front of embryonic outgrowths (Figure 3A). This STB contained densely packed nuclei and often occurred at multiple, distinct sites (Figure 3B). STB was also positive for human chorionic gonadotropin (hCG) subunit β (Figure 3C), which forms the maternal recognition of pregnancy hormone that is specifically secreted by STB from the onset of pregnancy.

**Figure 3.**
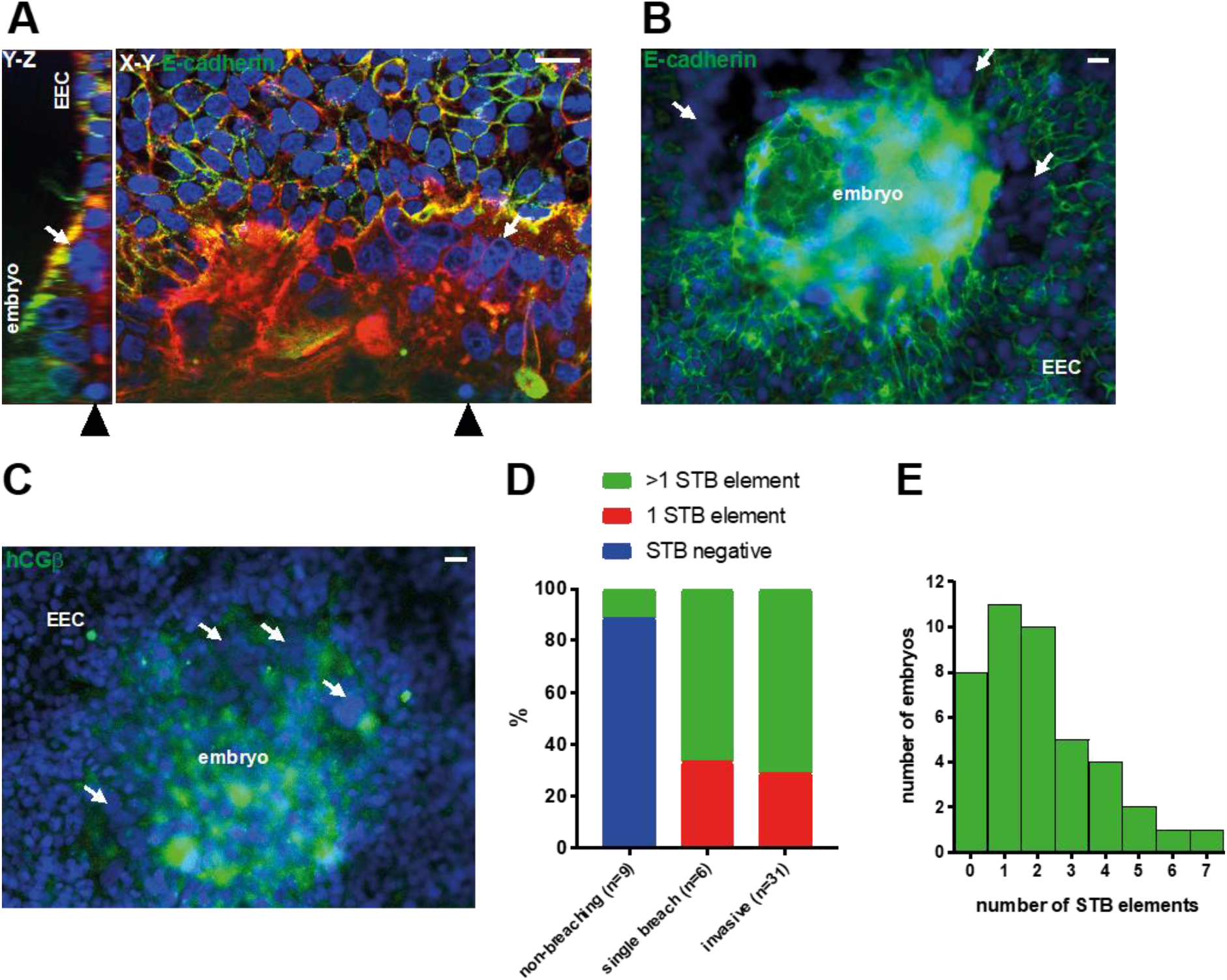
***A*** Fluorescence optical sections of an invasive embryo labelled with phalloidin (red), DAPI (blue) and anti-E-cadherin (green), panels show two X-Y planes. White arrows indicate STB. Scale bars 20μm. ***B*** Fluorescence micrograph of an invasive embryo labelled with DAPI (blue) and E-cadherin (green). White arrows point to distinct regions of STB. Scale bar 20μm. ***C*** Fluorescence microscopy image of an invasive embryo labelled with DAPI (blue) and anti-hCGβ (green). White arrows point to distinct regions of STB. Scale bar 20μm. ***D*** Bar graph relating STB quantities to embryo invasiveness (n=46). ***E*** Bar graph of STB quantities in embryos (n=42; the number of STB elements could not be discerned in four samples).

STB, as determined by combining DAPI with E-cadherin and/or phalloidin staining, was seen in only 1/9 non-invasive embryos (11.1%). All breaching and invasive embryos contained STB, with approximately two thirds containing more than one STB element (Figure 3D). Notably, STB was never observed in TE not in contact with EEC. Analysis of the number of STB elements (multinuclear structures bound by E-cadherin and/or phalloidin staining) present in each embryo demonstrated that up to seven could be found and >70% embryos had 1-4 distinct STB elements (Figure 3E).

### Ishikawa EEC layer stimulates STB differentiation in TSC spheroids

Human TSC can differentiate to form the major trophoblast subtypes, including STB (Okae *et al*. 2018), and thus offer a model with greater experimental scope than single embryos for interrogating trophoblast differentiation at implantation. Treatment of TSCs with the previously reported STB differentiation medium induced syncytialisation and upregulation of hCGβ at the mRNA and protein level (Supplementary Figure 1A, B). TSC culture in non-adherent conditions produced blastocyst-sized spheroids of mostly mononuclear cells (Figure 4A), which were co-cultured with Ishikawa EEC layers to model human embryo implantation *in vitro*. TSC spheroids attached to EEC layers within 6h (Figure 4B), reflecting the kinetics of blastocyst attachment (Ruane *et al*. 2020). After 48h, spheroids had invaded the epithelium with STB at the invasion front (Figure 4C). Invasive trophoblast was hCGβ-positive, although this was not specifically localised to STB (Figure 4D). The expression of STB markers in spheroids co-cultured with EEC layers and those cultured in isolation was compared to determine whether interaction with EEC induced STB differentiation. STB genes *CGB* and *SDC1* were significantly upregulated by co-culture with EEC layers (Figure 4E), while secreted hCGβ levels were also higher in co-cultured TSC spheroids (Figure 4F). These experiments suggest that EEC interactions stimulate STB differentiation from multipotent trophoblast.

**Figure 4.**
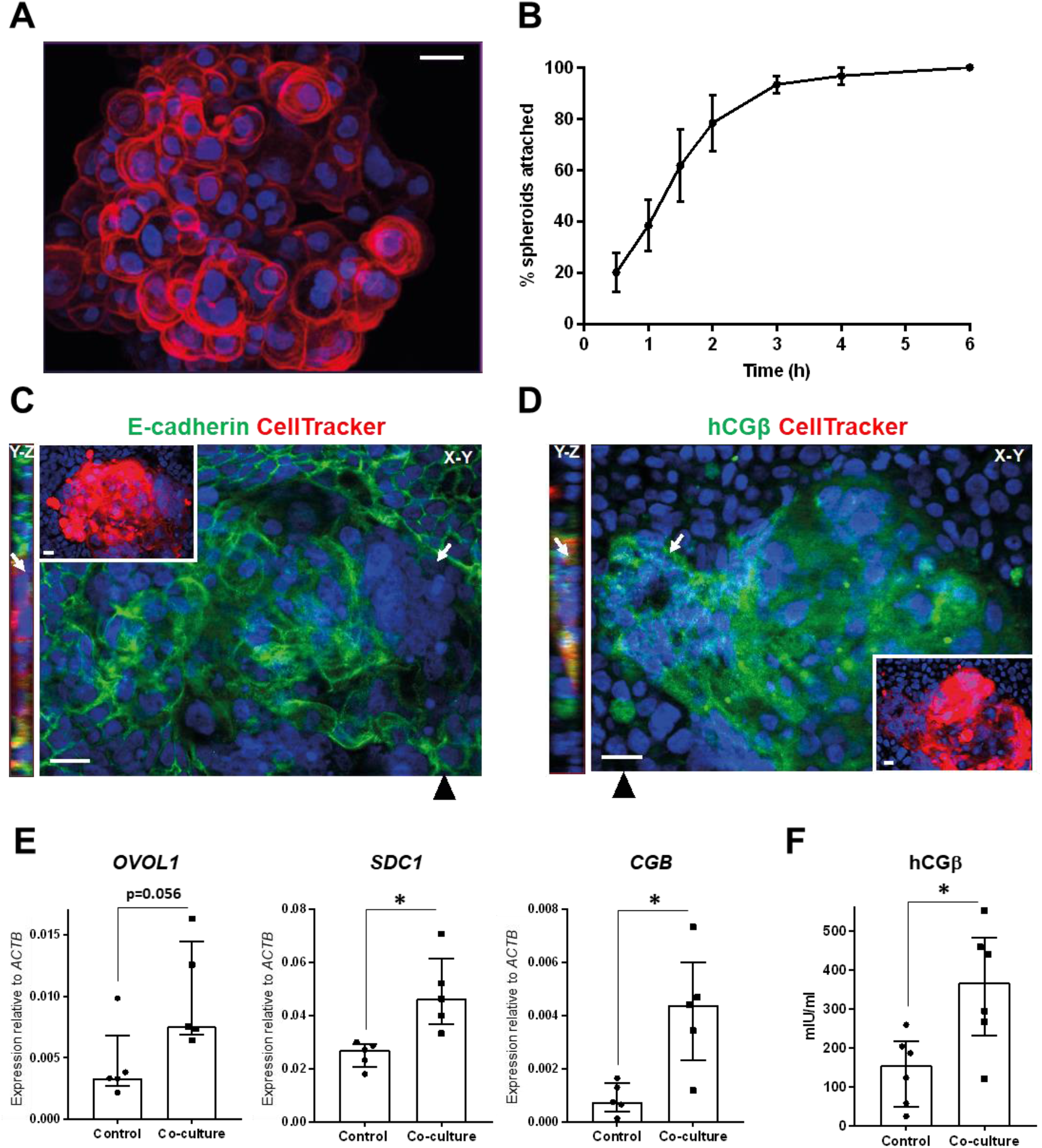
***A*** TSC spheroid labelled with phalloidin (red) and DAPI (blue), imaged by optical sectioning fluorescence microscopy and displayed as maximum intensity projection. Scale bar 20μm. ***B*** TSC spheroid attachment to Ishikawa EEC layers was monitored over 6h. Three independent repeats were performed (~20 spheroids per repeat), mean ± SEM plotted on line graph. ***C-D*** CellTracker-loaded TSC spheroids attached to Ishikawa EEC layers after 48h co-culture were labelled with DAPI (blue) and ***C*** anti-E-cadherin (green) or ***D*** anti-hCGβ (green), and optically sectioned by fluorescence microscopy. X-Y and Y-Z planes are shown as indicated, with black arrowheads indicating the location of the Y-Z plane. CellTracker (red) and DAPI labelling is shown in insets. STB is indicated by white arrows. Scale bars 20μm. ***E*** TSC spheroid-Ishikawa EEC co-cultures and isolated TSC and Ishikawa EEC cultures were lysed after 48h for qPCR analysis. Expression of STB markers *OVOL1, SDC1* and *CGB* was assessed relative to *ACTB* expression. Five experimental repeats, median ± IQR plotted, * p<0.05 Mann Whitney. ***F*** Medium was collected from TSC spheroid-Ishikawa EEC co-cultures and isolated TSC and Ishikawa EEC cultures after 48h for hCGβ ELISA. Six experimental repeats, median ± IQR plotted, * p<0.05 Mann Whitney.

### Gene networks at the TE-EEC interface

To identify the gene networks functioning at the TE-EEC interface, we used single-cell transcriptomic datasets from blastocyst TE (Petropoulos *et al*. 2016), and bulk transcriptomic data from EEC directly isolated from endometrium (Chi *et al*. 2020). Cell surface or secreted genes, differentially expressed in receptive (mid-secretory phase) *vs* non-receptive (proliferative phase) endometrial epithelium (Figure S2A), were used to identify 39 cognate cell surface genes in day 6 TE. The resulting *in silico* TE-EEC interface (Table 1), was enriched for extracellular matrix components, peptidases, cell adhesion molecule-binding proteins, signalling receptor-binding proteins and glycosaminoglycan-binding proteins (Figure S2B).

**Table 1.**
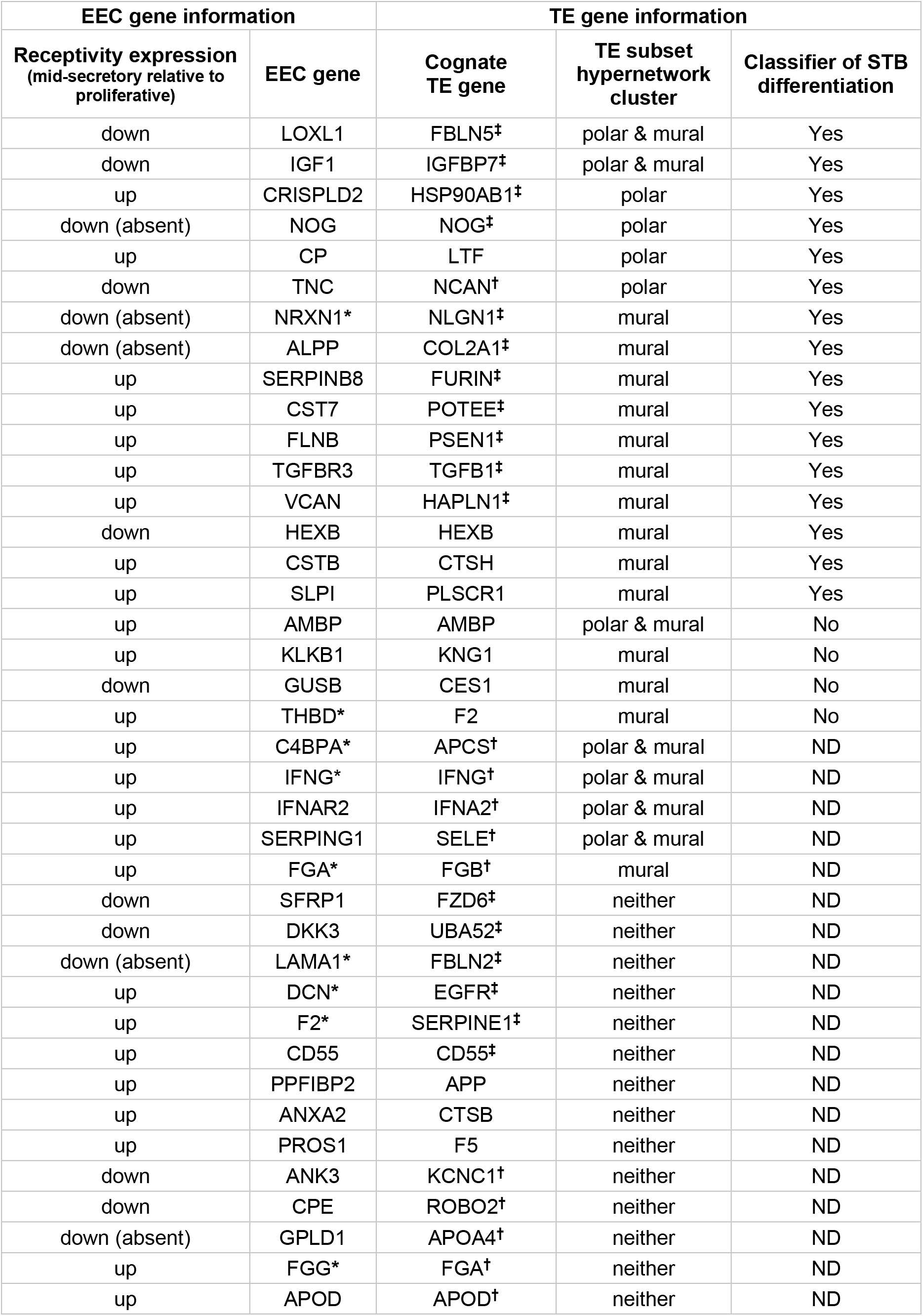
Summary of *in silico* TE-EEC interface, hypernetwork analysis and random forest classifications. * EEC gene not expressed in Ishikawa cells (RNAseq; data not shown); ^†^ TE gene not expressed in TSC; ^‡^ TE gene differentially expressed in TSC-STB differentiation; ND not determined.

Hypernetwork analysis enables clustering of genes based on co-expression within the transcriptome, which relates to connectivity in functional gene networks (Pearcy *et al*. 2016, Gaudelet *et al*. 2018). Day 6 TE subsets previously defined as polar and mural (Figure S2C) (Petropoulos *et al*. 2016), enabled us to use hypernetwork analysis to detect differences between polar and mural TE interactions with EEC. A cluster of 11 genes was identified from analysis of EEC-interacting polar TE genes (Figure 5A), while a 21-gene cluster was found from EEC-interacting mural TE genes (Figure 5B), with 7 genes common between the two clusters (Table 1).

**Figure 5.**
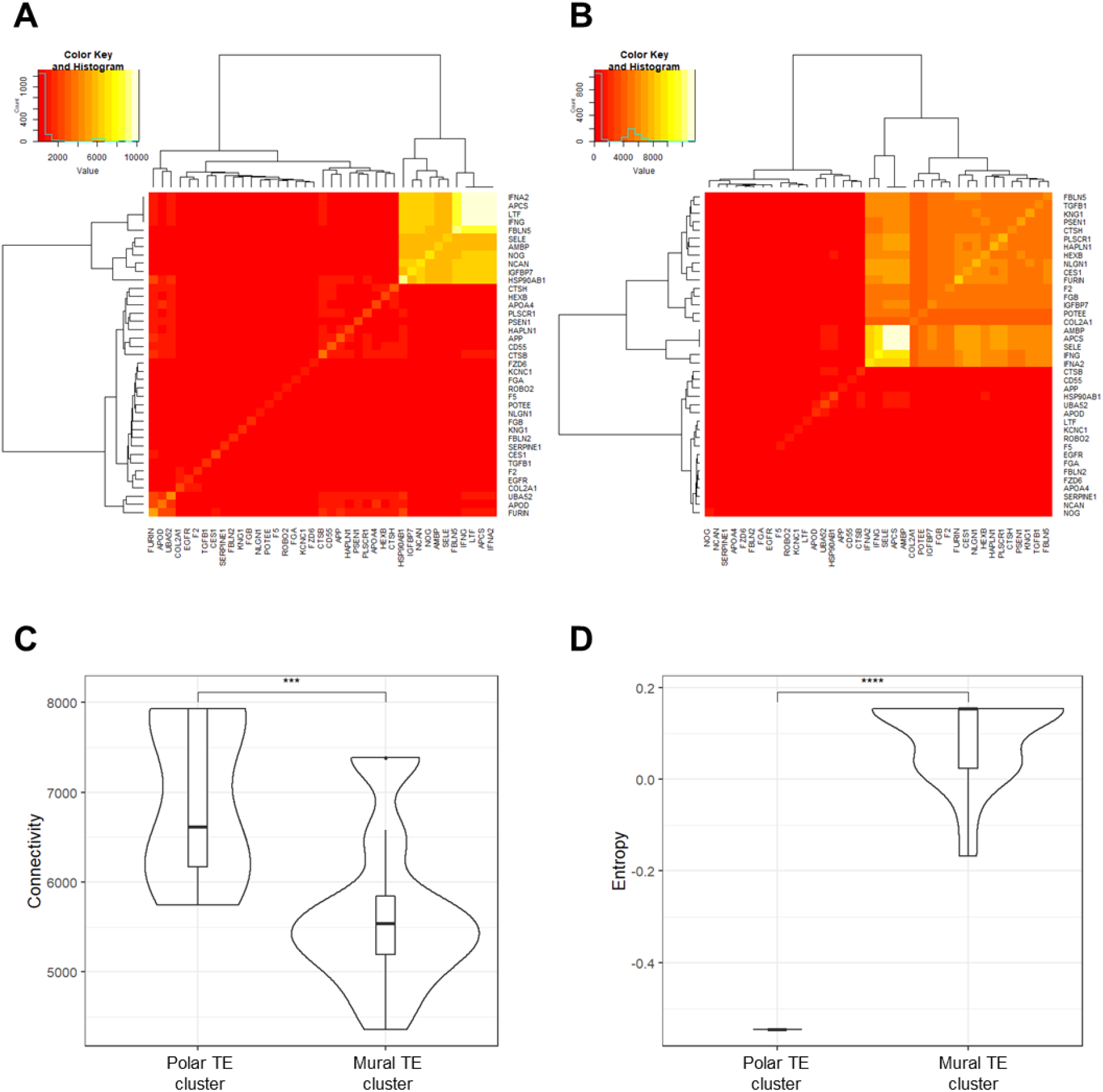
***A*** Hypernetwork analysis was performed on EEC-interacting TE genes (n=39) in polar TE cells (n=86) and presented as a heatmap. Eleven genes form the highly connected central cluster. ***B*** Hypernetwork heatmap of TE-EEC interface genes (n=39) in mural TE cells (n=245), with 21 genes forming the highly connected central cluster expanded. ***C*** Connectivity of hypernetwork clusters (the number of pairwise shared correlations) in polar and mural TE is presented as a violin plot. Median line and interquartile range is illustrated in each box, and whiskers illustrate the range with outliers as points. *** p<0.001 Wilcoxon rank sum test. ***D*** Entropy from the hypernetwork clusters of polar and mural TE (a measure of the organisation of gene connections), relative to entropy generated by permuting one thousand hypernetworks of random genes within the respective transcriptomes. Comparison of absolute entropy values between gene networks and subsets therein is not informative connectivity organisation. Data is presented in violin plot form as above. **** p<0.0001 Wilcoxon rank sum test.

The hypernetwork heatmaps suggested enhanced transcriptome connectivity in polar TE compared to mural TE. Quantification of the hypernetwork properties revealed a higher connectivity (number of shared transcriptome correlations) between EEC-interacting polar TE genes compared to those in mural TE (Figure 5C), and this reflected differences between background transcriptome connectivity in polar and mural TE (Figure S2D). Entropy is a measure of the organisation of gene connections within the hypernetwork, and is presented relative to background levels as comparisons of absolute entropy are not informative of connectivity organisation. A lower entropy, indicative of a more co-ordinated organisation of gene connectivity, was found for EEC-interacting polar TE genes than for those in mural TE (Figure 5D).

### In silico evidence for EEC-induced STB differentiation

As EEC interact with differentially connected gene networks in polar and mural TE in our model, we assessed gene ontologies within the hypernetworks (Table 2, 3). Genes connected to EEC interactors of both the polar and mural TE were over-represented in biological processes including chemical stimulus detection and innate immune responses. Notably, an enrichment of genes involved in G-protein-coupled receptor (GPCR)-coupled cyclic nucleotide signalling was found to be connected to both polar and mural TE genes. Activation of this pathway is known to induce STB differentiation (Knerr *et al*. 2005). To test whether the EEC interactors in polar and mural TE are linked to STB differentiation, we applied a random forest machine learning approach using genes from the hypernetwork clusters to investigate TSC differentiation to STB (Figure S3) (Okae *et al*. 2018). 7/11 polar TE genes were present in the TSC transcriptomes and 4 were differentially expressed upon differentiation to STB, while 16/21 mural TE genes were present of which 9 were differentially expressed. Strikingly, random forest analysis suggested 6/7 polar TE and 12/16 mural TE genes to be sufficient to classify STB differentiation from TSC (Figure 6A, B; Table 1).

**Figure 6.**
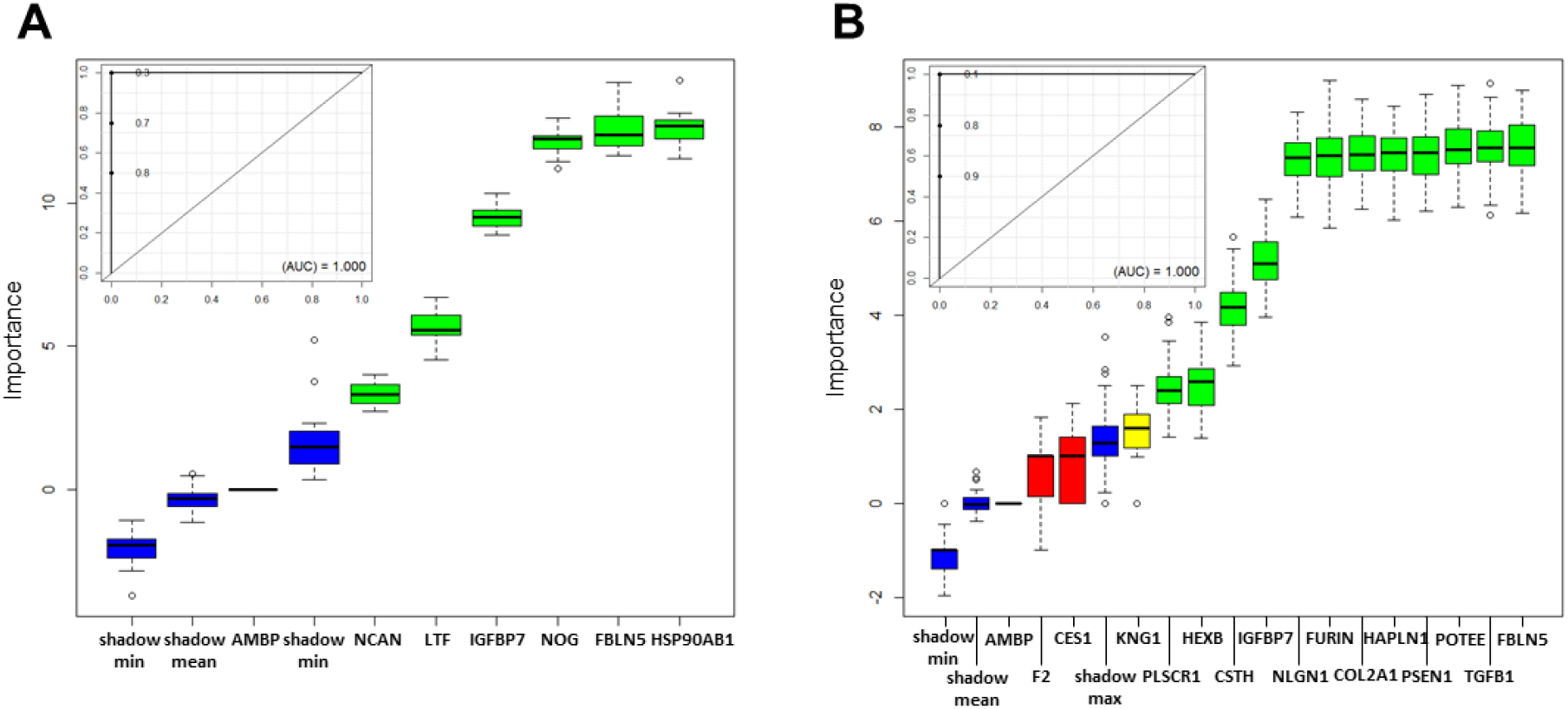
Boruta plots of random forest machine learning (1000 iterations) performed to test classification of TSC differentiation to STB (TSC n=4, STB n=4) for hypernetwork-identified EEC-interacting ***A*** polar TE (n=7) and ***B*** mural TE (n=16) gene networks, with corresponding ROC curves for informative genes inset. Box-whisker plots represent importance Z-scores, a metric of informativity of a variable for classification. Green confirms statistical significance in the classification of STB, red indicates a non-informative gene in the classification of STB, yellow indicates a failure to either confirm or reject a gene as informative within the allotted number of random forest runs, and blue demonstrates the background variation of predictive value within the data. Random forest determined that the hypernetwork-identified EEC-interacting TE genes confirmed in both polar (6/7) and mural TE (12/16) are informative enough to classify TSC from STB (area under ROC=1).

**Table 2.** Polar TE hypernetwork connectivity biological process gene ontologies.

| **Gene Set** | **Description** | **Size** | **No. of genes present** | **Enrichment Ratio** | **False discovery rate** |
| --- | --- | --- | --- | --- | --- |
| GO:0050906 | detection of stimulus involved in sensory perception | 478 | 275 | 6.44 | <1.00E-16 |
| GO:0007606 | sensory perception of chemical stimulus | 473 | 278 | 6.58 | <1.00E-16 |
| GO:0009593 | detection of chemical stimulus | 457 | 282 | 6.91 | <1.00E-16 |
| GO:0006959 | humoral immune response | 242 | 74 | 3.43 | <1.00E-16 |
| GO:0098542 | defense response to other organism | 473 | 97 | 2.30 | 4.91E-13 |
| GO:0007187 | G protein-coupled receptor signaling pathway, coupled to cyclic nucleotide second messenger | 250 | 64 | 2.87 | 6.13E-13 |
| GO:0018149 | peptide cross-linking | 60 | 27 | 5.04 | 2.13E-11 |
| GO:1990868 | response to chemokine | 93 | 33 | 3.97 | 1.38E-10 |
| GO:0015837 | amine transport | 101 | 29 | 3.22 | 7.53E-07 |
| GO:0001906 | cell killing | 157 | 40 | 2.71 | 7.53E-07 |

**Table 3.** Mural TE hypernetwork connectivity biological process gene ontologies.

| **Gene Set** | **Description** | **Size** | **No. of genes present** | **Enrichment Ratio** | **False discovery rate** |
| --- | --- | --- | --- | --- | --- |
| GO:0050906 | detection of stimulus involved in sensory perception | 478 | 212 | 8.06 | <1.00E-16 |
| GO:0007606 | sensory perception of chemical stimulus | 473 | 215 | 8.26 | <1.00E-16 |
| GO:0009593 | detection of chemical stimulus | 457 | 214 | 8.51 | <1.00E-16 |
| GO:0018149 | peptide cross-linking | 60 | 25 | 7.57 | 4.72E-14 |
| GO:0006959 | humoral immune response | 242 | 47 | 3.53 | 3.32E-12 |
| GO:0098542 | defense response to other organism | 473 | 70 | 2.69 | 3.32E-12 |
| GO:0030101 | natural killer cell activation | 85 | 20 | 4.28 | 2.94E-06 |
| GO:0007187 | G protein-coupled receptor signaling pathway, coupled to cyclic nucleotide second messenger | 250 | 37 | 2.69 | 3.77E-06 |
| GO:0007218 | neuropeptide signaling pathway | 101 | 21 | 3.78 | 1.02E-05 |
| GO:0097696 | STAT cascade | 148 | 26 | 3.19 | 1.09E-05 |

## Discussion

Placentation is essential for human development, and this process begins with blastocyst attachment to the endometrial epithelium and breaching of this barrier at implantation (Aplin & Ruane 2017). Here, *in vitro* and *in silico* models of the epithelial phase of human embryo implantation reveal for the first time that interactions with EEC induce TE differentiation to STB, which then goes on to breach the endometrial epithelium. Multiple STB elements were observed in *in vitro* implanting embryos and random forest-based classifications in an *in silico* model, which together indicate that EEC-induction of STB differentiation occurs in both polar and mural TE. Greater connectivity and organisation was found in EEC-interacting polar TE genes, perhaps reflecting preferential polar TE attachment to EEC. These findings establish the importance of EEC interactions to the development of trophoblast lineages at implantation. To further investigate the pattern of trophoblast development at implantation, sequential interactions with endometrial epithelium, subjacent stromal matrix and decidual cell types must be considered. Understanding these processes will illuminate infertility and obstetric disease mechanisms thought to be underpinned by defective trophoblast development at implantation (Aplin *et al*. 2020).

Blastocysts that had attached to Ishikawa EEC layers after 48h co-culture, from day 6-8, revealed a range of implantation stages, from those attached to the EEC apical surface to those that had extensively invaded into the epithelium. Blastocyst cavity collapse is consistently observed by day 8 in embryos cultured in isolation (Deglincerti *et al*. 2016, Shahbazi *et al*. 2016), and consistent with this, most embryos here displayed collapsed blastocyst cavities and invasive morphology. Blastocyst collapse and re-expansion is observed routinely in mammalian embryos in culture as a function of ion transport across an epithelial layer, and is considered to be physiologically normal (Biggers *et al*. 1988). STB was observed in all invasive embryos and consistently pioneered breaching of the epithelium, with these sites appearing to form between cells of the EEC layer. Both mononuclear trophoblast and STB were present at the leading edge of embryos invading laterally into the EEC layer, suggesting that the initial stromal phase of invasion may not be driven solely by STB (James *et al*. 2012, Knofler *et al*. 2019, Turco & Moffett 2020). However, the 2D nature of the *in vitro* EEC model and the absence of underlying matrix and stromal cell types may have influenced invasive cell behaviour. The well-differentiated endometrial adenocarcinoma-derived Ishikawa cell line provides a consistent, polarised, receptive 2D epithelial model that is difficult to generate using primary EEC (Campbell *et al*. 2000, Singh & Aplin 2015, Ruane *et al*. 2017), nevertheless 3D cultures of primary endometrial cells are required to better recapitulate implantation.

Our evidence suggests that STB differentiates from EEC-attached TE, often at multiple regions, before invading between the EEC. STB may negotiate invasion between EEC through intercellular junctional complexes between trophoblast and lateral EEC membranes that have been seen in ultrastructural studies in both human and monkeys (Enders *et al*. 1983, Smith *et al*. 1987, Bentin-Ley *et al*. 2000). The STB we observed in both implanting embryos and the TSC spheroid implantation model was highly invasive, and this primary STB contrasts with the canonical STB that forms from villous cytotrophoblasts and bounds the villi formed at later stages of placentation (Aplin 2010, James *et al*. 2012, Yabe *et al*. 2016). TSC spheroid interactions with Ishikawa cell layers induced STB differentiation, producing invasive STB morphologically very similar to that formed from embryos, thus leading us to favour a model in which the primary STB phenotype is induced by EEC. Formation directly from TE as opposed to from cytotrophoblast could also contribute to the unique phenotype of primary STB, and comparison between EEC-induced STB from TE and that from the more cytotrophoblast-like TSC (Dong *et al*. 2020) could provide evidence for this. EEC-induced invasive trophoblast differentiation from TE was previously shown in mouse blastocysts (Ruane *et al*. 2017), implicating evolutionary conservation of this mechanism at the onset of interstitial implantation.

We applied network biology approaches to model interactions at the TE-EEC interface, using bulk primary EEC and TSC transcriptomes, and single-cell TE transcriptomes. A prior network biology approach used whole embryo and whole endometrium microarray transcriptomes to identify key gene networks, including adhesion, extracellular matrix and cytokine genes (Altmae *et al*. 2012). While we found some of the same genes (APOD, VCAN, FBLN2, LAMA1, SERPINE1, TGFB1), our focus on EEC interactions with polar and mural TE subpopulations using RNAseq data provided a model with increased resolution. Hypernetwork analysis enabled enrichment of gene networks linked to TE function at implantation, and we have previously used this approach to investigate ICM and TE lineage allocation from the 8-cell blastomere stage in human embryos (Smith *et al*. 2019). The expression correlation matrices underlying hypernetworks capture gene networks that relate to functional units within the transcriptome, with the sensitivity of this correlation approach lending itself to highly dynamic biological systems (Pearcy *et al*. 2016, Gaudelet *et al*. 2018). In addition to this unbiased approach, a hypothesis-driven approach using random forest machine learning implicated a network of sixteen EEC-interacting TE genes as drivers of STB differentiation. Integration with our *in vitro* models revealed that these TE genes were all expressed in TSC, while expression of 14/16 mid-secretory phase primary EEC genes were matched in Ishikawa cells. Ishikawa EEC layers induced STB formation from embryos and TSC spheroids, suggesting that the STB in these *in vitro* models is comparable to that produced upon TE attachment to primary EEC.

Single cell transcriptomic discrimination of TE subpopulations have linked polar TE with upregulation of STB differentiation genes (Petropoulos *et al*. 2016, Lv *et al*. 2019). Here, we used these TE subpopulations to show increased levels and organisation of connectivity of EEC-interacting polar TE gene networks. Gene network properties of connectivity and entropy are indicative of cell differentiation state; low entropy characterises reduced regulatory cross-talk and pathway redundancy in more differentiated cells (Banerji *et al*. 2013, Teschendorff & Enver 2017). Our analysis therefore corroborates polar TE as more differentiated than mural TE, and also suggests that interactions with EEC are better linked to cell differentiation gene networks in polar TE than in mural TE. Polar TE-oriented attachment was observed here and in previous studies (Bentin-Ley *et al*. 2000, Deglincerti *et al*. 2016, Shahbazi *et al*. 2016, Aberkane *et al*. 2018), and embryonic pole-oriented stromal invasion was observed *in vivo* (Hertig *et al*. 1956). The more differentiated state of EEC-attached polar TE could lead to polar-oriented STB invasion. In contrast, we found both polar and mural TE connectivity to the GPCR-coupled cyclic nucleotide signalling pathway central to STB differentiation (Knerr *et al*. 2005), and random forest demonstrated a similarly high propensity for EEC-interacting polar and mural TE genes to drive STB differentiation. Moreover, we observed multiple STB regions in contact with Ishikawa cells. Together, our findings suggest a model whereby STB forms from EEC-attached polar and polar-proximal mural TE before initiating invasion into the endometrium. The corollary of this model is that distal mural TE not engaged with EEC gives rise to trophoblast stem cells. This scenario is different to that in mice, where mural TE initiates invasion and polar TE gives rise to multipotent trophoblast that go on to form the placenta (Sutherland 2003). Studies of differential phagocytosis activity between polar and mural TE support this inversion of embryonic pole functionality between mouse and human blastocysts (Rassoulzadegan *et al*. 2000, Li *et al*. 2016).

Further work with sophisticated *in vitro* models consisting of multiple maternal cell types are required to pursue some of the outcomes of this study and to fully characterise the establishment of trophoblast populations at implantation. Failure of this process is likely behind many of the recurrent implantation failures seen in assisted reproduction patients and may also contribute to cases of recurrent miscarriage (Koot *et al*. 2011). Furthermore, there is increasing awareness that later stage obstetric pathologies have their origin in implantation and early pregnancy (Rabaglino *et al*. 2015, Garrido-Gomez *et al*. 2017, Than *et al*. 2018, Aplin *et al*. 2020), and understanding endometrial shaping of trophoblast development is central to developing new treatments.

## Materials and methods

### Human embryos

Embryos generated for IVF treatment but not used by patients were obtained with informed written consent at Old Saint Marys Hospital, Manchester, or other IVF units in England.

Embryos from both current and previous (cryopreserved) treatment cycles were used in accordance with ethics approval from the NRES committee south central (Berkshire; REC reference: 12/SC/0649), and a research license from the Human Fertilisation and Embryology Authority (R0026; Person Responsible: Daniel Brison), centre 0067 (Old Saint Mary’s Hospital; fresh embryo research) and University of Manchester (0175; frozen-thawed embryo research). Day 6 blastocysts were artificially hatched using acid Tyrode (pH 2.5), as described (Ruane *et al*. 2020), before use. Four of the embryos were thawed from cryopreservation at pro-nuclear stage or on day 2 after fertilisation. The remaining 42 embryos were donated fresh from IVF cycles.

### Ishikawa cell culture

Ishikawa EEC (ECACC 99040201) were cultured at 37°C, 95% air and 5% CO_2_ in growth medium (1:1 Dulbecco’s modified Eagle’s medium:Ham’s-F12 (Sigma) containing 10% fetal bovine serum (Gibco) and supplemented with 2 mM L-glutamine, 100 ug/ml streptomycin and 100 IU/ml penicillin (Sigma)). Ishikawa cells were grown to confluency to form layers in 24-well plates (Greiner) on 13 mm glass coverslips for co-culture with blastocysts and TSC spheroids and subsequent microscopy, or in 48-well plates without coverslips for co-culture with TSC spheroids for subsequent gene expression and ELISA analysis.

### Human blastocyst implantation assay

Confluent Ishikawa EEC layers were washed and replenished with serum-free growth medium immediately prior to the transfer of hatched day 6 blastocysts to individual wells. After 48h, all co-cultures were washed in PBS and fixed with 4% PFA in PBS for 20 minutes.

### TSC culture and spheroid formation

TSC were acquired as a gift from Dr Hiroake Okae, and cultured and differentiated to adherent STB as described (Okae *et al*. 2018). TSC were seeded into Aggrewell™ plates (Stem Cell Technologies) at ~200 cells/microwell and cultured in serum- and BSA-free TSC medium for 48h to produce spheroids.

### TSC spheroid implantation assay

TSC spheroids were collected from Aggrewell™ plates and resuspended in co-culture medium (1:1 DMEM:Ham’s-F12, 1% ITS-X (Gibco), 5μM Y27632 (Adooq Bioscience), 2mM L-glutamine and 100 ug/ml streptomycin and 100 IU/ml penicillin). ~20 spheroids, pre-treated with CellTracker™ (Life Technologies), were co-cultured per well of Ishikawa cells in 24-well plates for 48h before fixation with 4% PFA. Attachment was monitored over the first 6h of co-culture by agitating the culture plate to determine which spheroids were attached to the Ishikawa EEC layer. Alternatively, ~400 spheroids per well of Ishikawa cells in 48-well plates were co-cultured for 48h, before lysis and RNA extraction.

### Staining and fluorescence imaging

Fixed samples were quenched with 50mM ammonium chloride solution before permeabilisation with 0.5% Triton-X100 PBS. Samples were incubated with primary antibody in PBS for 2h or overnight at 4°C, followed by alexa568-phalloidin (Life Technologies), 4’,6-diamidino-2-phenylindole (DAPI) (Sigma) and alexa488/649 secondary antibodies (Life Technologies) for 1h. Human embryo samples were mounted in a chamber of 3% 1,4-diazabicyclo[2.2.2]octane (DABCO) (Sigma) in PBS. TSC samples were mounted on a glass slide with 3% DABCO Mowiol (Sigma). Fluorescence microscopy was carried out using a Zeiss Axiophot microscope equipped with an Apotome module for optical sectioning. Images were processed using Zeiss Zen software and ImageJ. Antibodies: rabbit anti-E-cadherin (Epitomics), rabbit anti-CDX2 (Cell Signalling Technologies), mouse anti-GATA3 (R&D Systems), and mouse monoclonal anti-hCGβ 5H4-E2 (Abcam).

### RNA extraction and PCR

RNA was extracted using the RNeasy Mini Kit (Qiagen), according to the manufacturer’s instructions. Reverse transcription using random 9mer primers (Agilent) followed protocols for Sensiscript Reverse Transcription kit (Qiagen). Real-time quantitative PCR (qPCR) were performed using reverse transcription reactions together with 0.25μM gene-specific primers (*SDC1* CTGCCGCAAATTGTGGCTAC, TGAGCCGGAGAAGTTGTCAGA; *CGB* CAGCATCCTATCACCTCCTGGT, CTGGAACATCTCCATCCTTGGT; *OVOL1* TGAACCGCCACATGAAGTGTC, GACGTGTCTCTTGAGGTCGAA; *ACTB* AAGCCACCCCACTTCTCTCT, CTATCACCTCCCCTGTGTGG) and QuantiTect SYBR green PCR kit (Qiagen) on StepOne Plus machines; 40 cycles, 60°C annealing temperature. Raw data was analysed with StepOne Plus software to yield cycle threshold (Ct) values, which were expressed as 2^-ct^ relative to housekeeping gene, *ACTB*.

### ELISA

Media from TSC spheroid cultures and co-cultures was collected and subject to hCGβ ELISA using kit EIA-1496 (DRG Instruments GmbH), according to the manufacturer’s instructions.

### Modelling the TE-EEC interface in silico

All *in silico* modelling and analysis was performed in R version 3.4.2 (R Foundation for Statistical Computing). Differentially expressed genes (DEGs) were defined by ANOVA (cut-off p<0.01) between proliferative phase (n=4) and mid-secretory phase (n=4) endometrial epithelial RNA sequencing (RNAseq) transcriptomes (GEO Accession GSE132711) (Chi *et al*. 2020). Localisation of the protein products of these genes was investigated using the Database for Annotation, Visualisation and Integrated Discovery (DAVID) (Huang da *et al*. 2009a, Huang da *et al*. 2009b). Proteins defined as localised to any of ‘extracellular space’, ‘extracellular matrix’, ‘proteinaceous extracellular matrix’ or ‘cell surface’ were refined from the list of DEGs. Putative binding partners were identified using the Biological General Repository for Interaction Datasets (BioGRID) (Stark *et al*. 2006), and refined to those which were localised as above in day 6 TE single cell-RNAseq transcriptomes (n=331) (Array Express Accession E-MTAB-3929) (Petropoulos *et al*. 2016). Transcriptomic differences between samples were visualised by principal component analysis (PCA) and partial least-squares-discriminant analysis (PLS-DA). All gene ontologies were established using Webgestalt (Zhang *et al*. 2005).

### Hypernetwork analysis

Hypernetworks were used to quantify shared correlations between EEC-interacting TE surface genes across the rest of the transcriptome. Correlation matrices were generated using the day 6 TE single cell-RNAseq transcriptomes. All analyses were repeated for polar and mural TE separately, modifying the samples included in each experiment, using previously inferred TE subpopulations (Petropoulos *et al*. 2016). Correlation matrices were binarised using the distribution of r-values in order to include only the strongest present correlations. Correlations further from the mean r-value than ±1 standard deviation (SD) in each set were assigned ‘1’, whilst those between the SD boundaries was assigned ‘0’. Hypernetworks were generated by multiplying the binary matrix (*M*) by the transpose of that matrix (*M^t^*) to describe the number of shared correlations between a pair of TE genes. The central cluster of the hypernetworks, arranged using hierarchical clustering, represent a subset of EEC-interacting TE genes whose expression patterns are correlated with a large number of genes across the transcriptome. The connectivity (mean number of shared connections) and organisation (Shannon entropy) of the central clusters were quantified. These metrics were also calculated by permuting one thousand hypernetworks of random genes within the transcriptome.

### Classification of genes using random forest

EEC-interacting TE genes present in TSC and TSC-derived STB (Okae *et al*. 2018) were used for random forest machine learning (Breiman 2001), which was applied over one thousand iterations to test whether the genes could accurately classify STB differentiation. Noise within the random forest was modelled using Boruta (Kursa & Rudnicki 2010), and classification was judged using receiver operating characteristic (ROC) curves.

### Statistical comparisons

Statistical significance was measured by Chi-squared and Mann-Whitney analyses using Prism (GraphPad). ANOVA and Wilcoxon rank sum analyses were performed using R.

## Acknowledgements

We would like to thank the assisted reproduction patients who kindly donated their embryos to this research programme, and the clinic staff at the Department of Reproductive Medicine, St Mary’s Hospital, Manchester, UK; Manchester Fertility, Manchester, UK; and the Hewitt Centre, Liverpool Women’s Hospital, Liverpool, UK, who made this possible. We also thank Dr Hiroake Okae for the kind gift of the trophoblast stem cell line used in this study.

---

**Supplementary Figure 1.**
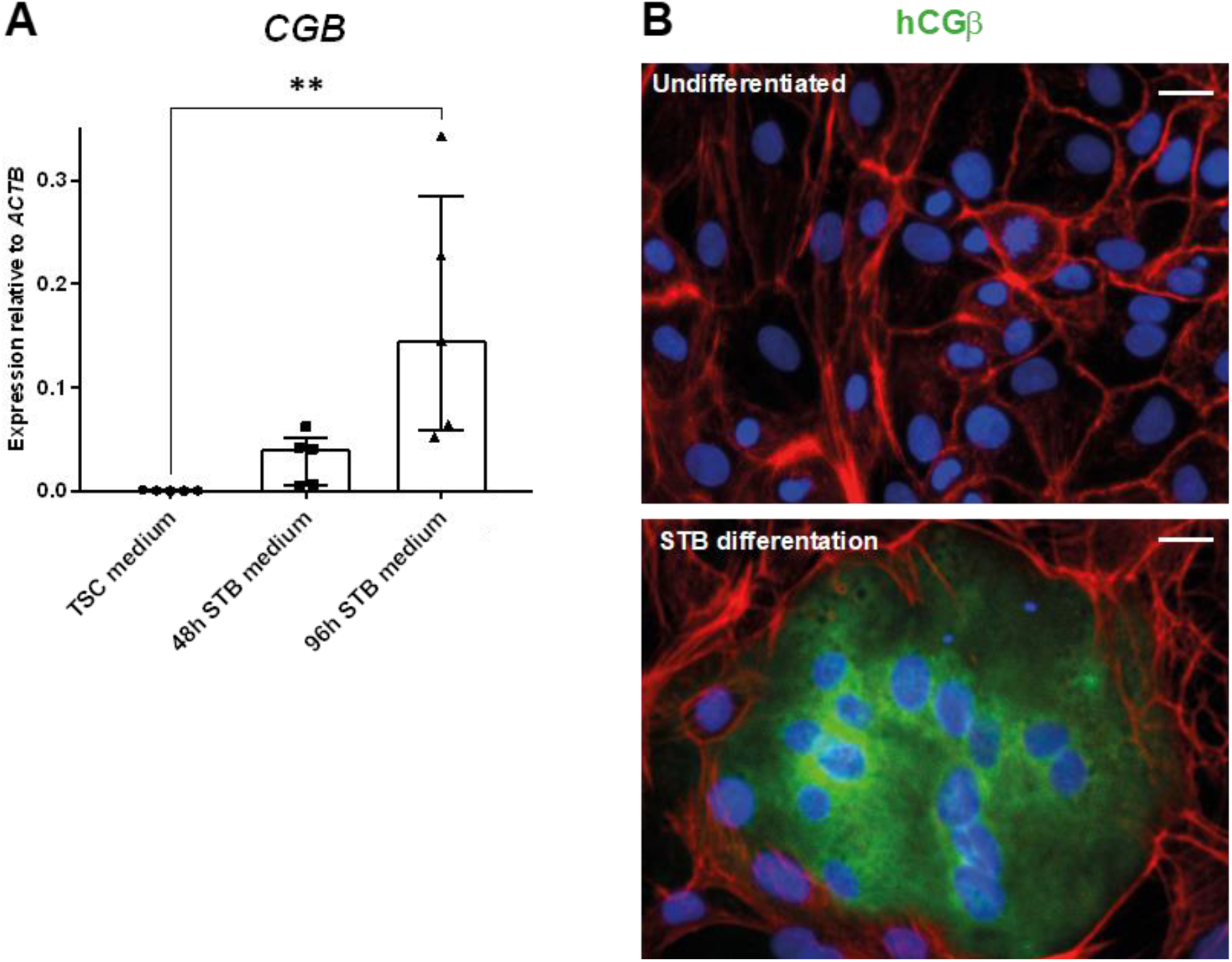
***A*** TSC were cultured in STB differentiation medium for 48h and 96h, and were compared to TSC cultured in TSC medium. Cells were lysed for qPCR and expression of STB marker *CGB* was expressed relative to *ACTB*. Five experimental repeats, median ± IQR plotted, ** p<0.01 Mann Whitney. ***B*** TSC were cultured in TSC medium and in STB differentiation medium for 96h. Cells were labelled with phalloidin (red), DAPI (blue), and anti-hCGβ (green). Scale bars 20μm.

**Supplementary Figure 2.**
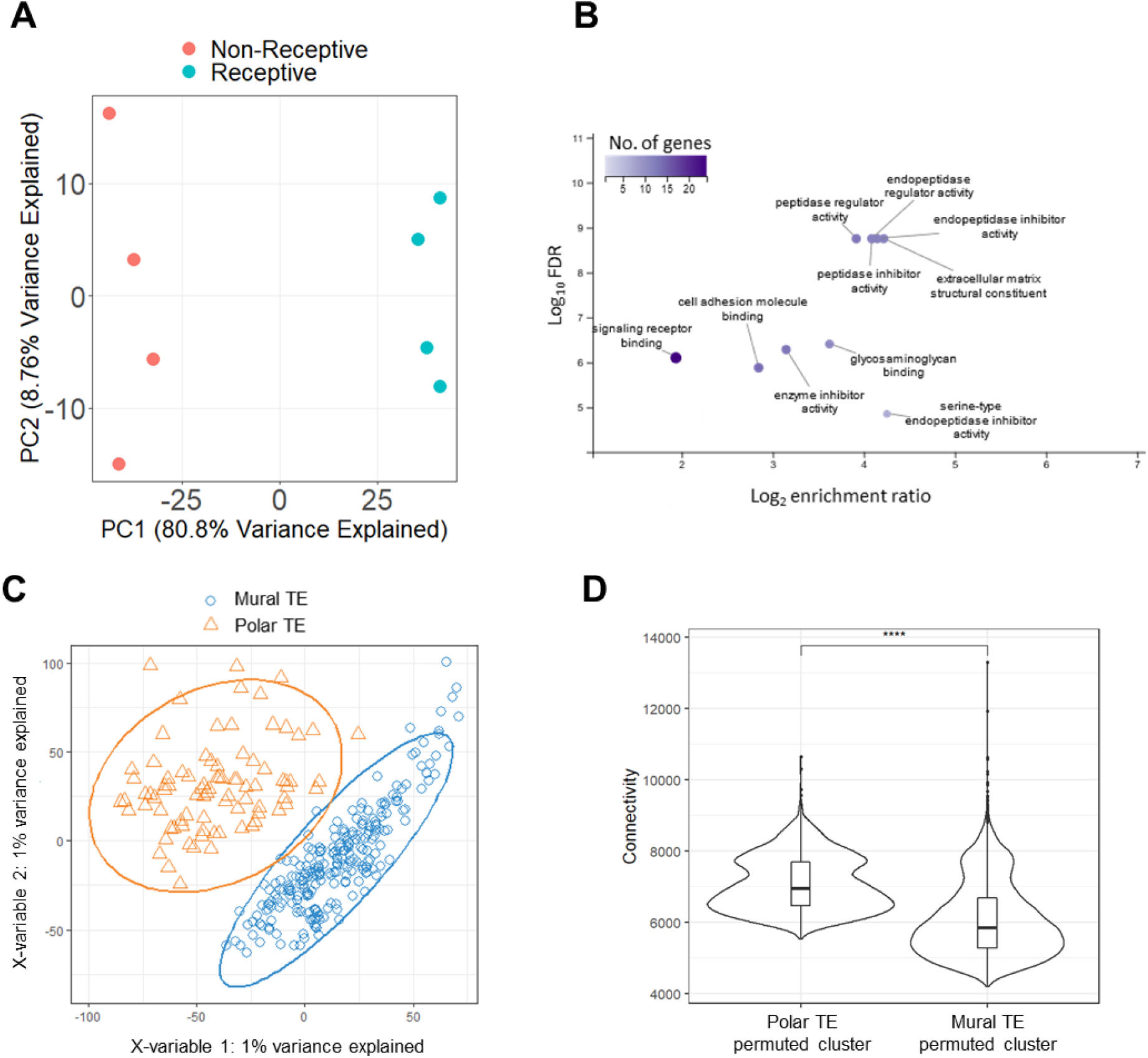
***A*** PCA of primary EEC transcriptomes from eight patients, four proliferative phase (receptive) samples (red) and four mid-secretory phase (non-receptive) samples (blue) (Chi *et al*. 2020). PCA calculated on DEGs (n=2131, p<0.01-8.53E-8) between proliferative and mid-secretory samples. Genes that were downregulated in mid-secretory EEC were not omitted in order to prevent introducing directional bias to the gene networks identified downstream. ***B*** Molecular function gene ontologies for TE-EEC interface genes, presented as false discovery rate (FDR) relative to gene-molecular function enrichment ratio. ***C*** Partial least squares-discriminant analysis (PLS-DA) of polar and mural TE (Petropoulos *et al*. 2016) performed on 331 whole TE transcriptomes (n=142780), demonstrating separation of TE into polar and mural based on a small fraction of the transcriptomes (Petropoulos *et al*. 2016, Lv *et al*. 2019). ***D*** Violin plots of background levels of polar and mural TE gene connectivity, as measured by permuting one thousand hypernetworks of random genes within the transcriptomes. Median line and interquartile range is illustrated in each box while whiskers illustrate the range, with outliers as points. *** p<0.001 Wilcoxon rank sum test.

**Supplementary Figure 3.**
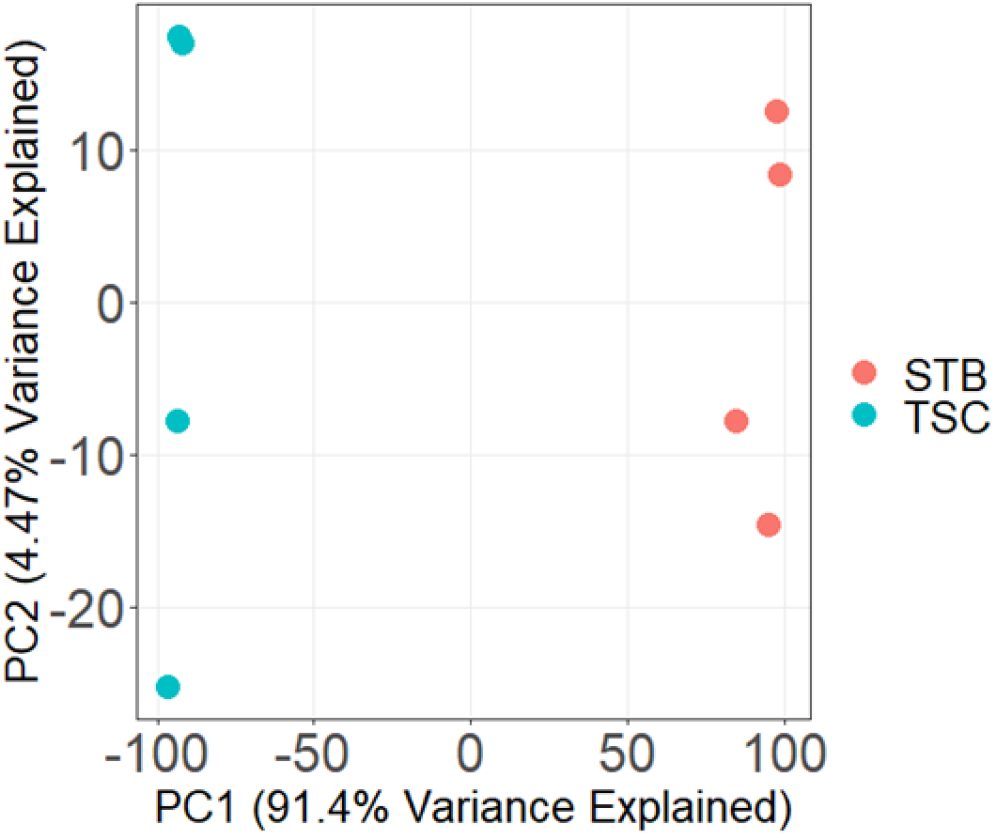
PCA of TSC (blue; n=4) and TSC-derived STB (red; n=4) (Okae *et al*. 2018). PCA calculated on whole transcriptome (n=24103).

